# Who removes seeds in a heavily urbanized area?

**DOI:** 10.1101/2022.12.21.521511

**Authors:** Dayana Hueso-Olaya, Maya Rocha-Ortega, Helí Coronel-Arellano, Fredy Palacino-Rodríguez, Alex Córdoba-Aguilar

## Abstract

In urban landscapes, granivores play a key ecosystem service: they contribute to the establishment of plants and thereby increase the probability of vegetal community recovery. It is therefore a priority to determine how and by whom seeds are removed, which seeds are removed and where they are removed. Our objective here was to determine the removal patterns in three core zones of an urban reserve, three urban parks, and three peri-urban parks, all in and around Mexico City. We performed a selective exclusion experiment with treatments for each type of seed consumer (ants, birds, and rodents) and seeds of nine plant species. Ants were the most important granivores in terms of seed removal (especially small seeds), followed by rodents and birds. Seeds of the native plants *Passiflora subpeltata, Pittocaulon praecox* and *Opuntia joconostle* had higher removal rates than other exotic and cosmopolitan species. The urban reserve had higher rates of seed predation compared to the urban and peri-urban parks. Thus, ants are pivotal in keeping seed removal and vegetation communities. We propose a series of measures to promote the ecosystem function of seed removal and increase plant diversity in these different urban patches.

## INTRODUCTION

The occurrence of native plants inside green areas in cities play critical roles in determining the quality and abundance of resources for wildlife that delivering ecosystem services [1]. Mostly in urban reserves conserve native species plants [2], while parks, corridors, and other types of urban green infrastructure more often contain cosmopolitan [3] and/or introduced species [4]. Besides, the spatial distribution of green infrastructure is relevant for determining ecosystem services [5]. Peri-urban areas and larger area are more biodiverse than smaller and inner-city areas due to lower human disturbance on natural vegetation [6,7].

Urban green infrastructure design is usually planned from an economic perspective, exploring social, and spatial attributes [8,9]. However, can also be directed to achieve specific goals, such as ecosystem services that are processes through which ecosystems and species support and enrich human life [10]. Urban ecosystem services (UES) concept has a yet unrealized potential to act as an object because needs to be carefully contextualized [11,12]. We conceptualized UES as a key species providing ecosystem functions and which their communities assembly and disassembly by urban environmental factors across spatiotemporal scales, lately translated in socio-economic factors [10–12].

Seed dispersers have been classified as mobile link species, which support essential ecosystem functions by moving actively or passively among green patches and ecosystems and overcome barriers [13]. Animal-mediated seed dispersal interactions may be particularly susceptible to the effects of urbanization negative affecting diversity and identity of the seed disperser [14], seed removal and behavior of seed dispersers due to high energetic cost of dispersion across the urban landscape [15]. Since economic point of view, average replacement cost of planting in urban parks was USD 22,500 in 2005 in Sweden, whereas for city natural oak forest regeneration was USD 9400 per hectare [13]. These estimates must motivate investments in management strategies that help secondary seed dispersers will contribute to plant establishment and regeneration of forest patches in human-landscape [16]. Instance of secondary seed dispersed are birds, insects, and mammals. Birds and insects are the most sensitive to human disturbance in terms of diversity, while mammals are in terms of interaction rates [17].

In this study, we evaluated the processes of removal of seeds across reserves and parks in an urban landscape in Mexico City (CDMX). Thus, our specific objectives were to: a) compare the patterns of seed removal among urban reserves (embedded in the city), urban parks (embedded in the city), and peri-urban parks (adjacent to the city); b) determine which species of plants are most removed, and c) identify taxonomic groups of granivores. We hypothesized that urban reserve due to their management will have higher rate of seed removal than peri-urban areas and urban parks [18]. Second, the introduced and cosmopolitan species will have the lowest rates of seed removal, because native species tend to have well established interactions in contrast to introduced species [4,19,20]. Given the type of vegetation (xerophytic scrub), we hypothesized that the most important granivores for removal will be the invertebrates (i.e., ants) [21].

## METHODS

### Study area

Arid and semi-arid regions occupy between 50 and 60 % of the total surface of Mexico. However, xerophytic scrub is the third most transformed ecosystem in the country [22] and the location of the largest cities, particularly Mexico city (CDMX), one of the most populous cities in the world [23]. In CDMX, the xerophytic scrub found on the lava fields of the Xitle volcano is a unique plant community due to the altitude at which it is found (between 2200 and 2500 m elevation). The zone has an area of 1495 km^2^, temperate rainy climate with maximum temperatures between 28 and 30 °C, mean annual precipitation below 250 mm and mean altitude of 2240 m. The type of vegetation is open scrub where the herbaceous stratum predominates and fills in during the rainy season [24], known as the Pedregal ecosystem. The flora includes approximately 377 taxa of vascular plants, of which 105 (28 %) are Asteraceae and slightly less than 10 % are introduced species [25]. This high richness of native vegetation, however, has been practically destroyed by the urban development [26].

### Urban green spaces

The sampling points were categorized to evaluate the effect of urban pressure on urban green spaces. For this categorization, we considered (I) management and purpose of the park, (II) structure of the vegetation present, (III) accessibility to people, and (IV) location with respect to the center of the city. Thus, this study had three categories: (1) Urban reserve—the Pedregal de San Ángel Ecological Reserve at the National Autonomous University of Mexico (hereafter, REPSA, for its acronym in Spanish), with an area of 1.77 km^2^ [27]. The REPSA is an ecological reserve embedded within the city that is dedicated to the preservation of xerophytic scrub and contains all of the plant strata associated with xerophytic scrub, a large number of herbaceous plants, few trees, and dominance of *Pittocaulon praecox* (Asteraceae). This zone was sampled in three polygons within the core zone, where human access is restricted (www.repsa.unam.mx). (2) Peri-urban areas at the city’s periphery were represented by two fragments of Bosque de Tlalpan National Park and Fuentes Brotantes de Tlalpan National Park. These two parks combine conservation and recreation, so some corridors within them are open to public use. Both areas contain mosaics of xerophytic scrub with introduced pine, cypress, and eucalyptus species (http://cdmatorral). (3) Urban parks, such as the Huayamilpas Ecological Park, Loreto and Peña Pobre Ecological Park, and Viveros de Coyoacán National Park. These areas are mainly dedicated to public recreation and are completely accessible to people. These areas do not have vegetative strata and there are few or no native species (http://red.ilce.edu.mx; http://cdmxtravel.com). All these urban parks are embedded within the city (Fig. 1).

**Figure 1.**
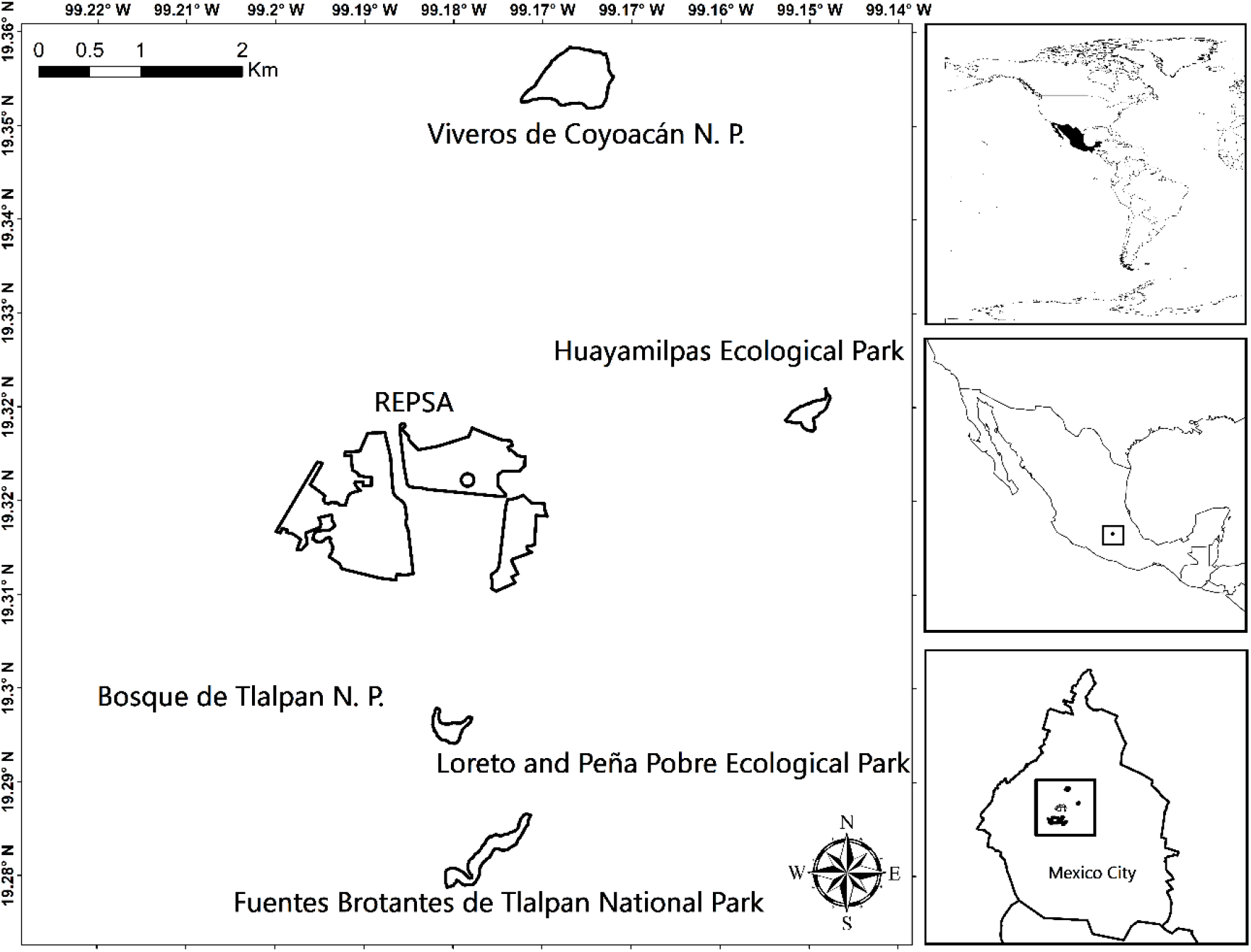
Location of urban green spaces used in this study in Mexico City, Mexico.

### Selection of seeds

The seeds used in the removal experiment were selected based on their biogeographic origin and distribution of the plant species. They were collected from the REPSA and the UNAM Botanical Garden. The seeds were categorized into the following three groups: (1) species endemic to Mexican xerophytic scrub and with restricted distribution, such as *Pittocaulon praecox* (cav.), endemic to the Pedregal ecosystem [24], *Dahlia coccinea* cav. is endemic to Mexico, and also the flower is representative of Mexico (http://www.conabio.gob.mx), and *Opuntia joconostle* Weber, endemic to central Mexico and are widely used in Mexican cuisine (http://www.conabio.gob.mx); (2) species native to Mexico and with widespread distribution, such as *Senna multiglandulosa* (Jacq.) (http://biologia.fciencias.unam.mx), *Passiflora subpeltata* Ortega (http://www.conabio.gob.mx), and *Manfreda scabra* (Ortega) (http://www.cm.colpos.mx); and (3) cosmopolitan species, such as *Dodonaea viscosa* (L.) (http://www.conabio.gob.mx), *Phytolacca icosandra* L. (http://www.conabio.gob.mx), and *Schinus molle* L. an introduced species (http://biologia.fciencias.unam.mx). All these species are common in the REPSA and fruit at the end of the dry season and beginning of the rainy season, which is the peak of activity for both invertebrates and vertebrates.

### Experimental design

Patterns of seed post-dispersal were evaluated using a series of selective exclusion treatments to identify the taxonomic groups responsible for the removal of different species of seeds. The exclosures treatments is a standard protocol widely used and effective to evaluated the granivores contributions to the seed removal [16,21,28–33], with easier control of overestimation of frequency of seed removal than visitation data [34]. The treatments were: (1) access to invertebrates, mainly ants, by inverting 2 oz cylindrical plastic jars over the seeds and fixing them to the ground, leaving a 1 cm^2^ opening, (2) access to ant and small rodents, placing 500 mL cylindrical plastic containers over the seeds, with a 6 × 6 cm opening, (3) seeds uncovered and accessible to invertebrates, rodents, and birds, and (4) no access (negative control to distinguish animal removal from secondary removal by air and rain) covering the seeds with 2 oz cylindrical plastic jars fixed to the ground [16,21]. In each urban green space, we established two 40 m-long parallel transects separated by 20 m. Along each transect, we placed five sampling stations, spaced every 10 m; at each station, we placed ten seeds for each of the four treatments, for a total of 40 seeds, for each of the plant species mentioned above. At each station, the treatments were placed a minimum of 50 cm apart from each other. The seeds of all nine species were offered simultaneously and were left exposed for 48 h, after which they were collected, counted, and weighed after being dried [16].

### Data analysis

We calculated the percentages of seed removed exclusive per each group (i.e., ant, rodent, and birds), subtracting the number the seed removed into treatment 1 to treatment 2, and the number of seed removed in both treatment 1 and 2 to treatment 3 to even the chance of seed consumption between treatments. Next, to evaluate the effects of exclusion treatments and type of urban green space (REPSA, peri-urban parks, and urban parks) on the rate of seed removal, we constructed independent generalized linear models (GLM) with a binomial error distribution for each of the seed species. The removal rate was the response variable and the treatment and urban green space were the explicative variables, along with their interaction. In the cases where the interaction was not significant, we repeat the analysis without including the interaction [35]. After each model, we did a post hoc to test the differences between level of variables. For all of the analyses we used R [36], and functions from the emmeans and stats packages [37]. Mean ± STD are provided unless stated otherwise.

## RESULTS

A total of 34.03 % of the seeds were removed (n = 3, 240). In general, the analysis of urban green space category showed that the REPSA had the highest rate of seed removal (5.19/10 seeds ± 3.16) followed by peri-urban parks (4.81/10 seeds ± 2.91) and finally, the urban parks (3.42/10 seeds ± 3.17). From the total seeds removed, 39 % were removed in the REPSA, 36 % in the peri-urban parks, and 25 % in the urban parks.

The species with the highest seed removal were *P. subpeltata* (5.61/10 seeds ± 4.1), *P. praecox* (4.67/10 seeds ± 3.3), and *O. joconostle* (4.05/10 seeds ± 3.7), followed by *D. viscosa* (3.47/10 seeds ± 3.2), *D. coccinea* (3.25/10 seeds ± 3.1), *P. icosandra* (3.02/10 seeds ± 3.5), *M. scabra* (2.66 ± 2.6), *S. molle* (2.3/10 seeds ± 2.3) and *S. multiglandulosa* (1.5/10 seeds ± 1.9). The experiments classifying seeds by their biogeographic origin and distribution showed that 39.91 % of native seeds with restricted distributions were removed, mainly of the species *P. praecox* and *O. joconostle*; followed by 32.59 % of widely distributed native species, with the species *M. scabra* and *S. multiglandulosa*. For the introduced and/or cosmopolitan seeds, the removal value was 29.59 %, mainly the species *D. viscosa* and *P. icosandra*.

The highest number of seeds was removed from treatment 3 (i.e., ants, rodents, and birds; 5.02/10 seeds ± 3.08), followed by treatment 1 (i.e., ants; 4.20/10 seeds ± 3.26), and finally, treatment 2 (i.e., ants and rodents; 4.20/10 seeds ± 3.24 SD). Treatment 4 (negative control) showed that a minimal number of seeds are removed by wind or rain (0.18/10 seeds ± 1.19). We found that from the total of seeds removed, 78 % were removed from treatment 1 (i.e., ants), 7.8 % of the seeds were removed exclusively by rodents, and 14.2 % were removed exclusively by birds.

The independent models for each plant species showed a differential effect of exclusion treatment and type of urban green space on the rates of seed removal for all plant species (Table 1). For *O. joconostle, P. subpeltata, D. viscosa*, and *P. icosandra* treatment, type of green space, and their interaction affected the rate of seed removal (Table 1). For *O. joconostle and P. icosandra*, treatment 1 had the highest removal rate in the REPSA. For *P. subpeltata*, treatment 2 had a highest removal rate in peri-urban parks. For *D. viscosa*, treatments 2 and 3 had highest removal rates in REPSA (Supplementary material 1).

**Table 1.**
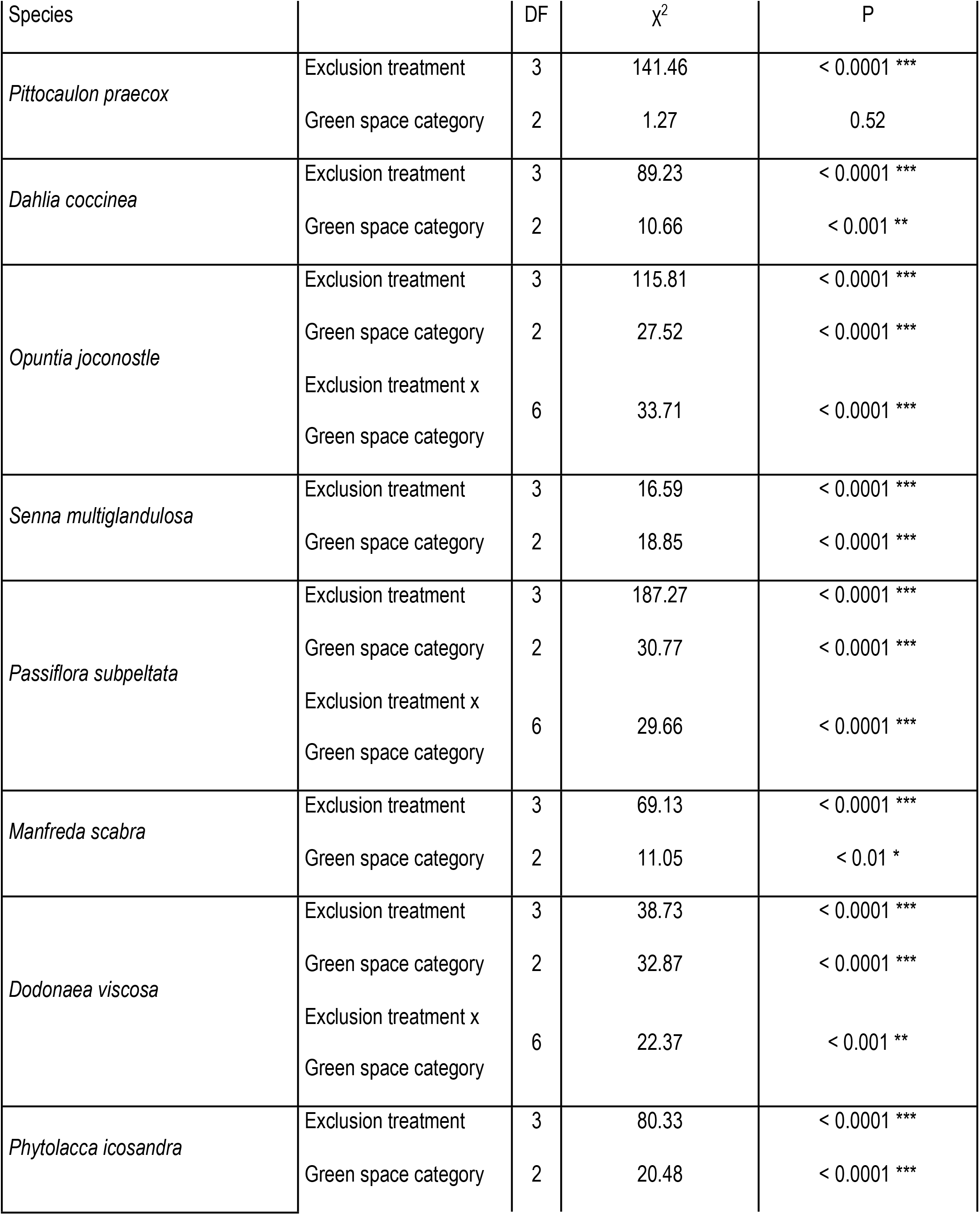
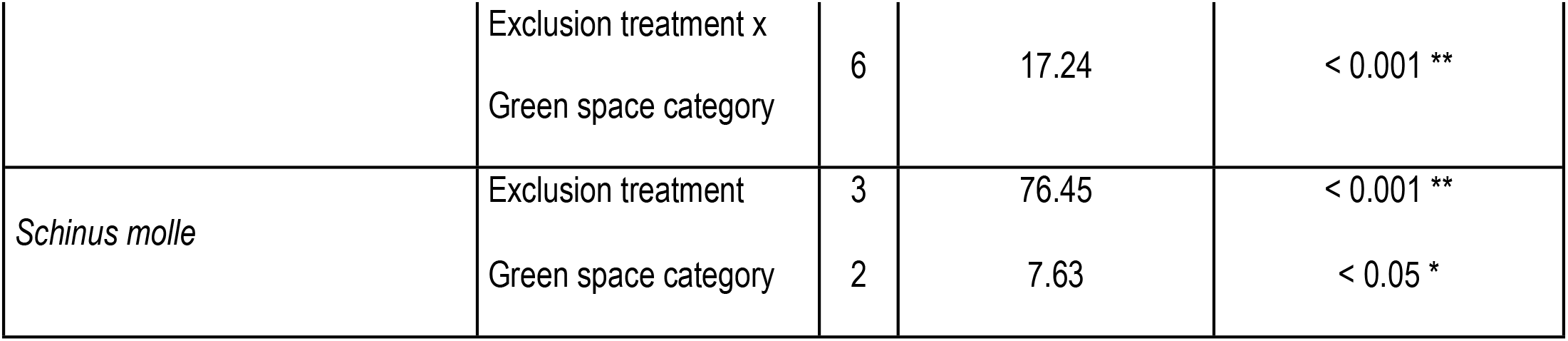
Effects on seed removal rate of each green space category (urban reserve, peri-urban park, urban park) and granivore exclusion treatment (invertebrates, small mammals, birds).

For *D. coccinea, M. scabra, S. molle*, and *S. multiglandulosa*, both treatment and the type of urban green space affected removal rate, but the interaction between these variables was not significant (Table 1). The highest removal rates for *D. coccinea* and *S. molle* were in the REPSA and peri-urban parks, while for *M. scabra* was in the REPSA. For *S. multiglandulosa* was greatest in treatment 3 in peri-urban parks, for *D. coccinea and S. molle* was in the treatment 3; finally for *S. molle* was similar among treatments (Supplementary material 2 and 3).

For *P. praecox*, only the exclusion treatment had an effect on seed removal (Table 1), with treatment 1 having a higher removal rate (Supplementary material 2).

In the REPSA, the seed removal rate was highest for the species *D. coccinea, O. joconostle, P. subpeltata, M. scabra, D. viscosa, P. icosandra* and *S. molle*. In peri-urban parks, the highest removal rates were for *S. multiglandulosa*, and *P. praecox* did not differ in removal rate among the different urban green spaces.

Treatment 1 (ants) had the highest seed removal rate for all of the plant species, particularly for *P. praecox*. In treatment 2 (ants and rodents), the highest removal rate was for *O. joconostle, P. subpeltata, D. viscosa* and *S. molle*. In treatment 3 (ants, rodents, and birds), the maximum seed removal rate was for *D. coccinea, S. multiglandulosa, P. subpeltata, M. scabra* and *S. molle*.

## DISCUSSION

According to our hypotheses, we found that native species with restricted distributions had higher rates of removal than introduced or cosmopolitans, particularly *P. praecox* and *O. joconostle*. These native disturbance-adapted species promote microenvironments for the establishment of shade plant species, production of aerial biomass, and nutrient exchange; all of these functions favor the restoration of the original vegetation [38]. If they germinate successfully, these plants may help to re-establish native fauna by increasing the availability of feeding and refuge sites [39]. We found intermediate removal rates for cosmopolitan species, such as *D. viscosa* and *P. icosandra*, but the lowest rate of removal was to introduce specie *S. mole*. Cities need to be understood within the context of their species interactions [40], because the disruption of interactions between legitimate seed dispersers (mutualist) with plants, beside by the introduction of exotic or cosmopolitan plant species [41] can lead to in the long term to homogeneous species assemblages and local extinctions [34,42]. Overall, our results suggest that both mutualistic– antagonist (i.e., seed disperser and predator) interactions still working in green areas of the city, particularly the interaction with native plants, likely because native species tend to established stronger interactions than introduced species [43].

We found that the removal of 78 % of the seeds of the 9 species was by ants, which demonstrates the important role that these organisms play in the seeds’ post-dispersal in an urban landscape. Ant assemblages shows a continuum in resilience to human disturbance, that is, some species are more resilient than other [44]. Furthermore, ants work as an important local filter that strongly suppress plant exotic recruitment due to are smaller than native [45]. In our study, ants mainly removed lighter native seeds of *S. praecox, P. icosandra* and *M. scabra* which are the lighter seeds of the scrub, therefore ant are the main post-dispersers of both native and cosmopolite species in the Pedregal ecosystem which emphasizes their role in the conservation of urban environment. On the other hand, the heavier seeds (i.e., *O. joconostle, P. subpeltata, D. viscosa* and *S. molle*) were removed by rodents, which removed 7.8 % of seeds and birds removed 14.7 % of the seeds, respectively. In general, vertebrates play an important role as secondary seed post-dispersers of heavier seeds in urban landscapes. However, fragmentation and urbanization tend to have negative effects on wild rodents and birds [46,47]. Birds are the most sensitive to anthropogenic disturbance, in terms of diversity, while mammals are negatively affected in their rates of interaction [17]. For both groups, the urban matrix may represent a negative selective pressure on movement among green patches [48]. To alleviate this, several studies have proposed corridors for vertebrates to move among urban green spaces by arranging them as a network [18,49]. This network would connect green spaces as “stepping stones” that would allow species to “hop” between green spaces.

We observed that rodents played a relatively minor role in seed removal, contrary to findings in more pristine arid environments [21]. The reason for this may be that some specialist granivore rodents have been locally extirpated, as is the case of the mouse *Lyomis irroratus* (Gray, 1868) [50–53]. Additionally, we speculate that native rodents could be displaced by exotic species such as *Rattus rattus* (Linnaeus, 1758), and *Mus musculus* Linnaeus, 1758. Overall, small-mammals could be more negatively affected than ants or birds by extirpation and introduction of species caused by urbanization.

Future studies are needed to evaluate the fate of seeds to discriminate between seeds that are dispersed or predated [54]. Moreover, although we were able to identify the taxonomic groups of granivores that participate in the post-dispersal of seeds, we were not able to identify the particular species of granivores, therefore, more studies would clarify not just this issue, but also the origin, distribution, and natural history of granivores [34]. Thus, for example, it is necessary to evaluate the contribution of exotic fauna to seed removal, since introduced species like the house mouse (*Mus musculus* Linnaeus, 1758) and black rat *Rattus rattus* (Linnaeus, 1758) that develop easily in urbanized environments could function as seed predator [55].

Complementary analyses such as the study of interactions between native and exotic fauna to understand both primary and secondary seed dispersion also are important [39,56]. In particular, we propose evaluations of the impact of cats on the abundance, density, and interactions of small and medium-sized native mammals [56]. In fact, cats could be responsible for the extirpation of specialist granivore rodents (e.g. *Lyomis irroratus*) [57]. In addition, is necessary to evaluate the contribution of vertebrate species in the process of primary removal in scrublands, as the mammal *Bassariscus astutus* (Lichtenstein, 1830), which is a very abundant frugivorous species in CDMX [58,59] and their interactions with plant species that have fleshy fruits (e.g., *P. icosandra*) that could increase the possibility that will colonize new environments (Ness et al., 2016). Fill these gaps, would provide a more complete picture of seed dispersion by the different actors and their threats inside the city.

## CONCLUSIONS

We found that conserved green areas, with the original plant structure, with a high proportion of native plant species and little access by humans, are sustainable regardless of their location within the city and maintain several of the natural ecosystem services such as secondary seed removal. We suggest three actions to the active restoration of the original vegetation of the urban green areas, first the use of seeds or cuttings to the reforestation with native plants in “exclusion” zones where human access is restricted to favor seedling reestablishment. This action might contribute to the establishment of native granivore communities that could contribute to the passive restoration of the vegetation at a low economic cost. Second, the eradication of feral fauna, mainly cats, to promoting the conservation of wildlife communities. Finally, control cosmopolitan and introduced plant species in the urban green spaces with scrub vegetation via remotion. Effective management of urban green spaces will allow the conservation of native flora and fauna [60], as well as interspecies interactions, even in small fragments embedded in the city.

In particular, we suggest that native plant species such as *P. praecox* and *O. joconostle* be used for active restoration. *S. multiglandulosa, D. viscosa*, and *M. scabra* could also be used in both active and passive reforestation projects, particularly in erosion soil, although with caution [61]. For its part, *D. coccinea* could be used as a landscaping element in gardens, corridors, and medians. In the case of *P. subpeltata* and *P. icosandra*, are an important source of food for frugivores and granivores, therefore must be favor their presence in green spaces, although *P. icosandra* should be control its, due to cosmopolitan distribution. Finally, it is advisable to control species such as *S. molle* given that is introduced species in the Pedregal ecosystem and being an arboreal element that is not natural in the ecosystem.

## Acknowledgements

This work was supported by PAPIIT projects IN206618 and IN204921. We thank all the attention brought by the staff of REPSA, Municipality of Coyoacan, Bosque de Tlalpan National Park, and Loreto and Peña Pobre Ecological Park. Also, we thank Grupo de Investigación en Odonatos y otros artrópodos de Colombia (GINOCO), Grupo de Investigación en Biología (GRIB) of the Biology Department at Universidad El Bosque, and the Centro de Investigación en Acarología.

